# Overestimation in time reproduction: Influences of accuracy feedback and attentional sharing

**DOI:** 10.1101/2023.08.22.554259

**Authors:** Lingyue Chen, Zhuanghua Shi

## Abstract

Duration reproduction is often subjective to biases, with a general tendency to overestimate durations, which has been observed in many studies. Yet, this overestimation is frequently dismissed as a nuisance error, and its underlying mechanisms remain elusive. Here, we conducted two experiments to investigate this general overestimation in duration reproduction. To pin down the origin of the error, we manipulated the reproduction output through shortened visual feedback in Experiment 1, while varying the presence of accuracy feedback in Experiment 2. Across both experiments, we observed a consistent overestimation in reproduction when accuracy feedback was absent. This overestimation, amounting to approximately 13.5% on a ratio basis across different durations and sessions, was unaffected by shortened visual feedback. We propose that this consistent overestimation is likely due to the attentional sharing between the action execution and the monitoring of the passage of time during the reproduction process.

## Introduction

In everyday life, we are immersed in a sea of temporal information. How we perceive, process, and integrate this temporal information is fundamental. For a trivial task, such as catching a flying ball, we need to constantly estimate the approaching time and adjust our own response. Researchers of time perception have developed various timing tasks, such as duration reproduction, temporal discrimination, and categorization (including temporal bisection or generalization), to facilitate this understanding.

An interesting phenomenon is that we generate estimation errors across various durations of which we are not explicitly aware. Two types of biases have been identified in time estimation. The first, commonly known as the central tendency bias, has been well-documented 150 years ago (Glasauer & Shi, 2021; Hollingworth, 1910; Lejeune & Wearden, 2009). The central tendency bias is exhibited when participants are asked to judge a set of durations. Shorter durations are often overestimated and long durations underestimated. This type of bias has been extensively studied, with a typical explanation being that we implicitly construct a prior centered around the set of durations. This prior is then integrated with the current duration (Bausenhart et al., 2014; Cicchini et al., 2012; Glasauer & Shi, 2021; Jazayeri & Shadlen, 2010; Shi, Church, et al., 2013; Shi & Burr, 2016), causing judgments to gravitate toward a mean duration. Varying the spacing or distribution of sample intervals can shift the temporal bisection point, a middle point of judgments between the given ‘short ’ and ‘long ’ standards, toward the ensemble mean (Baykan et al., 2023; Zhu et al., 2021).

The second type of bias, often referred to as a constant error or general bias, involves a general overestimation or underestimation in time perception (Craig, 1973; Grondin, 2001, 2012). For instance, intervals perceived as “filled” are often deemed longer than “unfilled” intervals (Craig, 1973), and auditory intervals are typically perceived longer than visual ones (Wearden, Goodson, et al., 2007). Reproducing a given auditory interval usually leads to an overestimation (Grondin, 2012; Shi, Ganzenmüller, et al., 2013). In a series of reproduction tasks where participants were asked to reproduce durations ranging from 1000 to 1900 ms, Grondin (2012) found consistent positive errors in reproducing intervals, except for the longest 1900 ms interval. Reproducing a one-second auditory interval with a simple button press could result in a 40% overestimation (Shi, Ganzenmüller, et al., 2013). Similar constant errors were also observed when participants reproduced a standard duration presented together with a comparison duration, particularly when the standard was presented second (Bausenhart et al., 2014). Although such general bias is common in duration judgments, it often garners less attention in interpretation and is treated as nuisance error and occasionally removed before further analysis (e.g., Cicchini et al., 2012). The prevailing explanation for this general bias is based on the internal-clock model. It postulates that either the speed of the clock varies (Wearden et al., 1998), or the timing of switch onset and offset differs (Craig, 1973).

In the case of the duration reproduction task, there are several factors that could contribute to a general bias. One noticeable difference between the duration reproduction task with other timing tasks is the involvement of action. It has been postulated that sensory modalities and the motor system may have distinct timing systems, and that these systems might process timing in a distributed manner (Bueti et al., 2008; Mauk & Buonomano, 2004). The timing of the motor system may be slower than the auditory or visual clocks, leading to consistent over-reproduction. A different, yet closely related, theory is the attentional sharing account (Fortin, 2003; Lejeune, 1998). When attention is shared between two subtasks, monitoring the passage of time and reproduction action, for instance, the limited attention resources could result in less attention being dedicated to monitoring the passage of time. Consequently, ‘ticks ’ may be lost, leading to an overestimation as reproduction time lengthens to compensate for the misstated counts. Contrasting the attention sharing account, attention might also switch between motor action and monitoring the passage of time (Zakay & Block, 1996). Attention could initially be focused on the action onset, then shift to time monitoring. When the duration approaches the target duration, attention would return to the action for stopping reproduction. Such attentional shifts can also cause general bias, differing from attentional sharing. Alternatively, it is also possible that attention remains on motor timing without switching to reproduction feedback, resulting in a general bias solely caused by the motor system. Notably, the attentional shift account predicts a constant error, independent of reproduced durations, while the attentional sharing theory suggests that the error would be proportional to the test duration.

On this premise, this study designed two experiments to investigate mechanisms underlying constant overestimation in the temporal reproduction task. One approach to distinguishing these alternative accounts involves manipulating temporal discrepancies between the action and action output. If reproduction relies solely on the motor timing system, the reproduced duration would base on the action itself, unaffected by variations in the action output (such as a delayed visual output). For instance, a shortened visual output wouldn ’t cause the reproduced duration to shrink. Conservely, if reproduction hinges on action output, adapting to a shortened visual output would shorten the reproduced duration. If attention is shared between action and action output, the reproduced duration would be overestimated with simultaneous action output according to the attentional sharing account, but would be partially shortened after adapting to a shortened action output. We tested this in Experiment 1. In Experiment 2, we varied test durations to discern the role of attention. This was achieved by introducing a short and a long duration and varying the presence of accuracy feedback. If attention switches between the two subtasks, the overestimated duration should remain constant for both short and long durations. Alternatively, the overestimated duration should vary when reproducing the short and long standard durations, maintaining a constant ratio to the reproduced duration.

## Experiment 1

### Method

#### Participants

A total of 20 volunteers, aged 19 to 31 (average of 23.55, 9 females and 11 males), were recruited for Experiment 1. Each participant had either normal or corrected-to-normal vision, and was naive to the purpose of the study. The sample size was determined by previous studies (Ganzenmüller, 2013), in which a group of 12 participants had yielded significant findings concerning under- and over-estimations. All participants signed the informed consent form prior to the experiment and received 9 Euros per hour for their participation. The experiment was approved by the Ethics Committee of the LMU Munich Faculty of Psychology and Pedagogics.

#### Apparatus

The experimental code, developed using the Psychopy (Peirce et al., 2019), controlled the presentation of visual stimuli on a ViewPixx LCD monitor (VPixx Technologies Inc.) with a refresh rate of 120 Hz. Behavioral responses were collected via a standard keyboard. The setup was housed in a sound-isolated dark cabin. The viewing distance was fixed to 60cm with the aid of a chin rest.

#### Stimuli and Procedure

A duration reproduction task was used in Experiment 1 (Figure 1A). A typical trial started with a central fixation cross, prompting participants to fixate on it. After 400 ms, a Gabar patch (size: 1.7° of visual angle, spatial frequency: 0.08 cycles per degree) appeared centrally for 800 ms, serving as the target duration, before disappearing. Participants then, with self-paced, attempted to replicate the target duration by holding down the spacebar. A Gabor patch, identical to the initial one, appeared as visual feedback during the key pressing.

**Figure 1.**
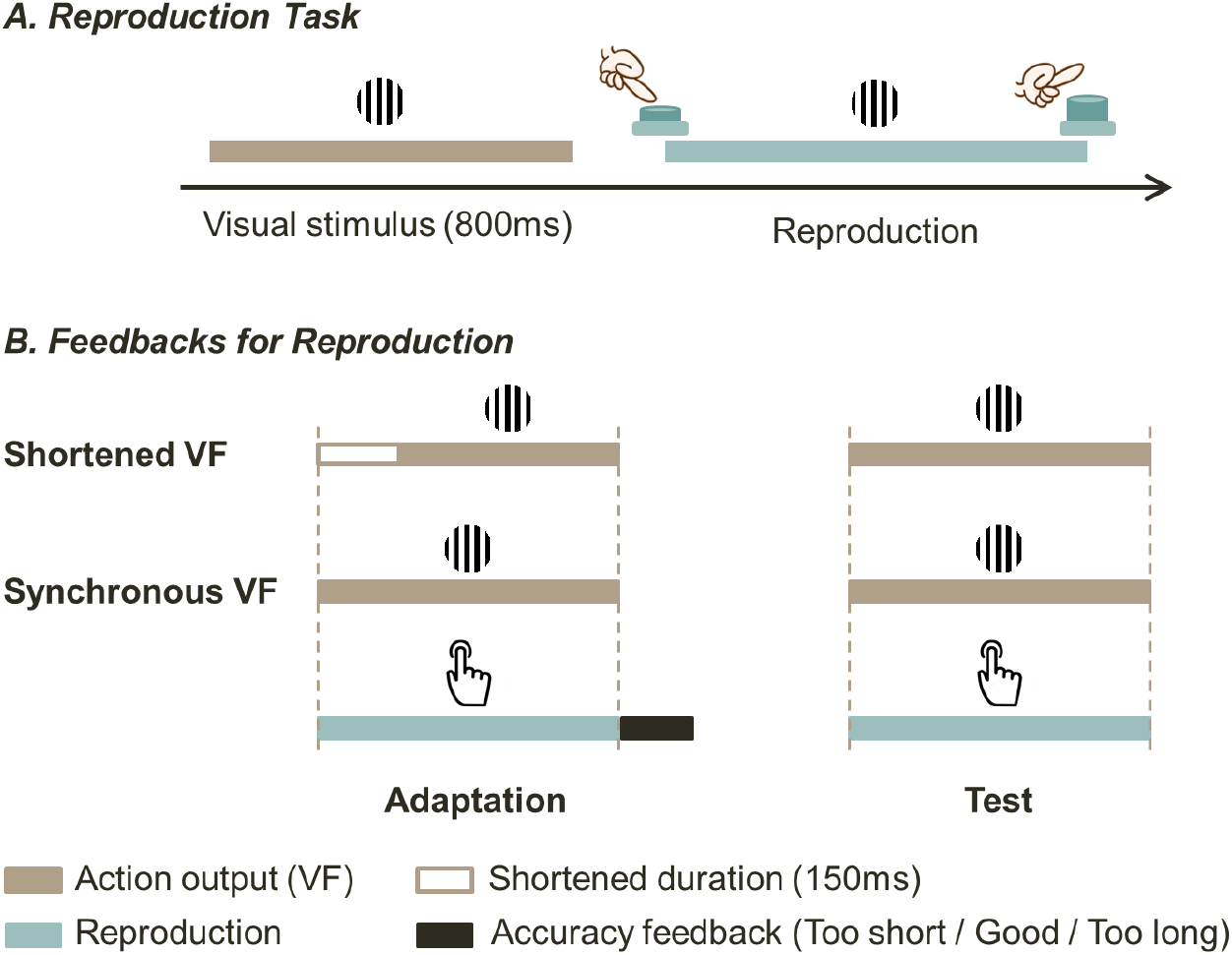
**Schematic illustration of the paradigm in** Experiment 1. The reproduction task **(A)** for each trial first presented a standard 800 ms target duration through a visual Gabor patch. After the visual stimulus disappeared, participants reproduced the target duration through pressing the spacebar. **(B)** Experiment 1 comprises two sessions, a shortened output adaptation (Shortened VF) and a simultaneous output adaptation (Synchronous VF). Each session comprised two phases: adaptation and test phases. In the session of the shortened output adaptation, the action output -visual feedback -started 150 ms later than the action onset, but stopped when the action stopped (left-upper panel). In the session of the simultaneous output adaptation, the reproduction was synchronized with the visual feedback. Upon the completion of the reproduction, an accuracy of the reproduced duration was provided (too short, long, or too long) based on the tolerant range of 10% error. The accuracy feedback was absent in the test phase.

The experiment comprised two sessions, namely the synchronous visual feedback (VF) session and a shortened VF session. Two sessions were tested successively with a 5 min break in between, and their order was counterbalanced across participants. Each session consisted of two phases: an Adaptation phase and a Test phase. Both sessions began with the adaptation phases (Figure 1B, left panel) with 60 obligatory trials. The key difference between two sessions was that in the shortened VF session, the visual feedback started 150 ms later than the action onset, but stopped when the action stopped, while the visual feedback synchronized with the action onset and offset in the synchronous VF session. Participants received immediate accuracy feedback based on how closely they replicated the target duration with motor action. The acceptable tolerance range was set at 10%, spanning from 720 to 880 ms), known as the ‘good ’ range. If the reproduced duration fell below this range, they received the text feedback ‘too short ’. Conversely, if the reproduced exceeded this range, the textual feedback ‘too long ’ was provided. Participants had to reach 80% of the last ten trials to the ‘good ’ range to finish the adaptation phase. If not, ten more trials were administered. 75% of participants managed to reach the goal within 60 trials, and 85% reached the goal with one round of ten more trials. Only one participant required 30 more trials to reach the goal.

Reproduction in both test phases employed the same visual stimuli with the synchronous visual feedback (Figure 1B, right panel). This phase consisted of 13 mini-blocks, with 20 trials each. The first five trials of the mini-block were top-up adaptation trials, each with accuracy feedback after the task, while the rest 15 trials provided no accuracy feedback. The target duration was consistent in all trials types (800ms).

### Data Analysis

The statistical analyses were performed using R (R Core Team, 2022). To ensure comparability across participants, only the final 60 trials of the Adaptation phase were included in the analysis, since some participants required more trials to meet the accuracy criterion. Additionally, the first trial of each sub-block in the test phase was excluded as they were influenced by the between-block break.

### Result and discussion

Experiment 1 investigated whether the shortened action-output (visual feedback) would affect the overestimation. During data preprocessing, one outlier was identified and excluded according to the interquartile range (IQR) rule. Figure 2B depicts the mean reproduced duration as a function of the mini-block trial number within the testing phase, encompassing the five top-up adaptation trials (brown lines) and 15 test trials (green lines). Two noteworthy patterns emerged during the test: a consistent lengthening in reproduction for the synchronous VF test trials relative to the shortened VF test trials, and an overall upward trend throughout the test phase, highlighting a distinct general bias characterized by growing overestimation across consecutive trials. Figure 2A details the mean reproduced durations for the Adaptation (the last 60 trials) and Test phase (excluding the top-up adaptation trials), separated by the Shortened-VF and Synchronous-VF sessions. The mean reproduced duration (± SE) were 788 (± 7), 893 (± 14), 809 (± 8), and 834 (± 19) ms for the Synchronous-VF/Adaptation, Synchronous-VF/Test, Shortened-VF/Adaptation, and Shortened-VF/Test conditions, respectively (Figure 2B).

**Figure 2.**
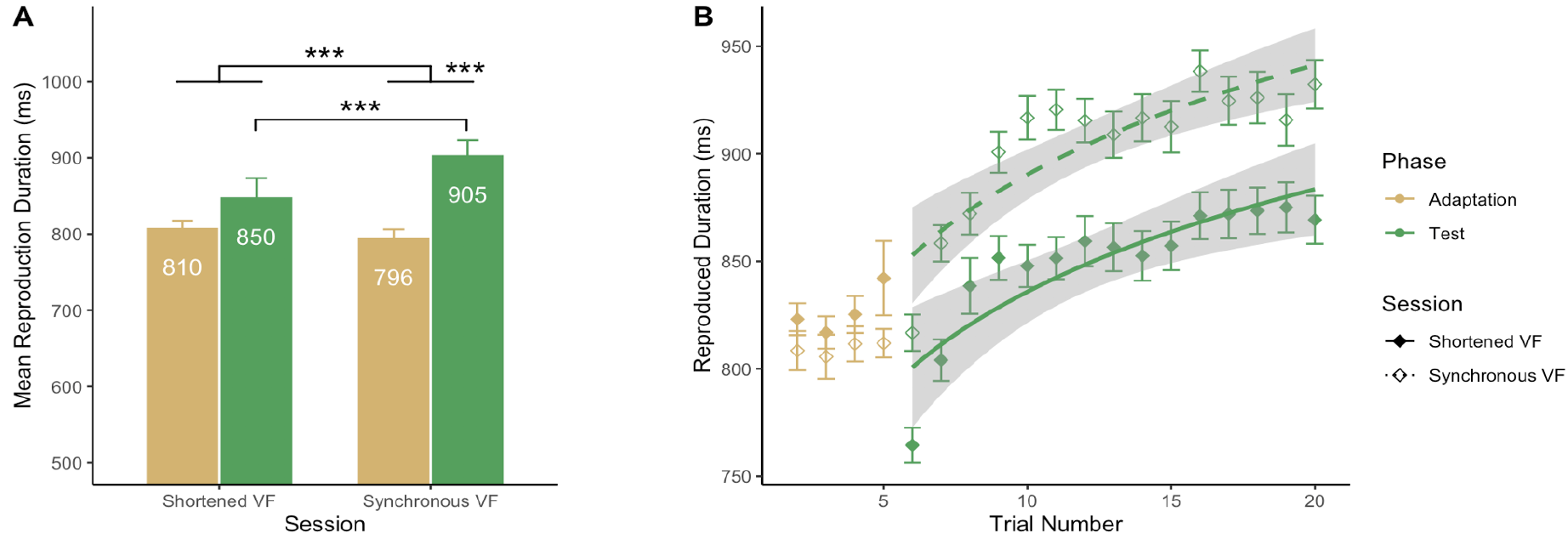
The general mean reproduction duration **(A)** for all four conditions (two sessions x two phases). The Trial-by-trial mean reproduction duration in the Test Phase **(B)**, comprising five top-up Adaptation trials and 15 Test trials in each sub-block. The fitted logarithmic regression model for the Test trials in both sessions **(B, right panel)**. The asterisks mark the significant interaction effect and between-condition differences (p < .001: * * *).

**Figure 3.**
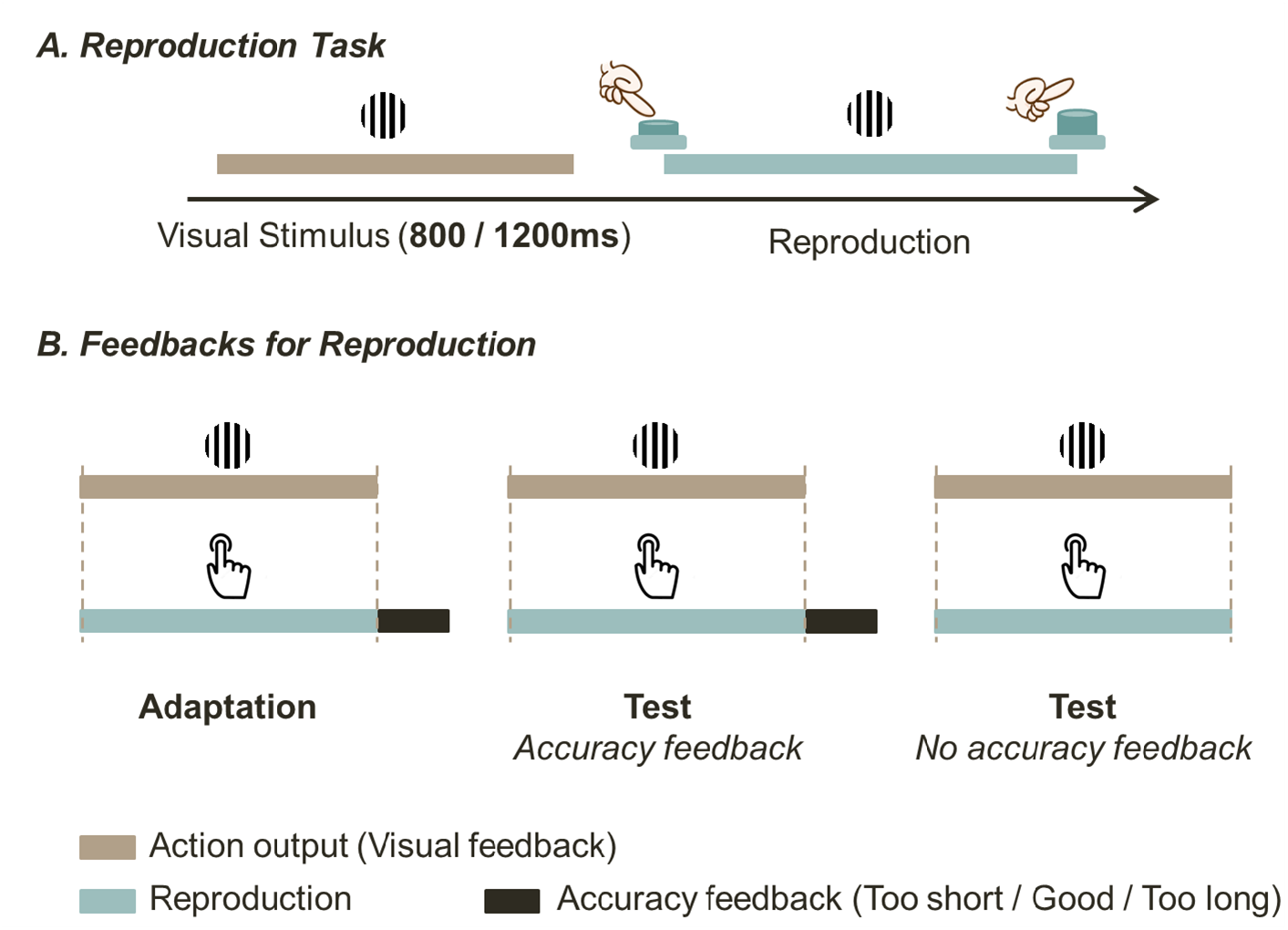
Schematic illustration of the paradigm in Experiment 2. The same reproduction task (**A**) was employed in Experiment 2 with two standard target durations, 800 ms and 1200 ms, tested in two separate sessions. Except for the standard duration, all settings in the two sessions were the same. The action output (visual feedback) was synchronized with the reproduction in both adaptation and test phases (**B**). Experiment 2 also applied the Adaptation-test paradigm. The test phase (middle and right panel) comprised four mini-blocks, two accuracy feedback and two no accuracy feedback blocks, presented alternatingly with an interleaved order. The accuracy feedback was only provided in the adaptation phase (left panel) and the accuracy feedback test blocks (middle panel).

A repeated-measure analysis of variance (ANOVA) with factors of Session and Phase revealed significant main effects for both Session (*F*(1, 18) = 6.60, *p* = .019, 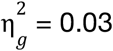), and Phase (*F*(1, 18) = 17.69, *p* < .001,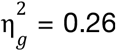). This demonstrated a significantly lengthened reproduced duration in the test phase than in the adaptation phase, as well as in the Synchronous-VF session than the Shortened-VF session. Moreover, the Session x Phase interaction was also significant (*F*(1, 18) = 28.82, *p* < .001, 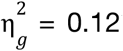). Post hoc pairwise comparisons revealed the interaction was mainly contributed by the longer reproduced duration during the synchronous-VF test phase (*t*s > 4.8, *p*s < .001), the comparable reproduced durations between the adaptations and Shortened-VF test phase (*t*s < 2.56, *p*s > .079).

We then further estimated the reproduction growing curve from the mini-test blocks with a logarithmic regression model (Figure 2B, right panel):

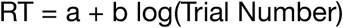

The curve was fitted separately to each session. Analysis of the logarithmic regression model revealed the following coefficient estimates for the two sessions. The mean intercepts were 673 and 729 ms for the shortened-VF and the synchronous-VF sessions, respectively. The former was significantly shorter than the latter (*t*(18) = 2.41, *p* = .027). The logarithmic slopes were comparable: 64.25 and 65.53 for the shortened-VF and synchronous-VF sessions, respectively. The slopes did not differ significantly (*t*(18) = 0.12, *p* = .908), indicating the growing overestimation was similar in two sessions in the absence of the accuracy feedback.

In short, Experiment 1 showed a significant influence of the shortened visual feedback on the duration reproduction in the test trials where accuracy feedback was not provided, suggesting that reproduction is not merely motor timing in the presence of action output. Rather reproduction is an integration of the motor and sensory feedback (Shi, Ganzenmüller, et al., 2013). The findings also demonstrated the effective updating of internal priors with accuracy feedback during the adaptation phase, as the accuracy feedback was based on purely motor action. However, this adaptation did not eradicate the general overestimation (absolute mean error of 109 ms) observed in the test phase when the accuracy feedback was omitted. In fact, reproduced durations gradually drifted back to overestimation, indicating reproduction is neither purely visual nor motoric timing. However, there remains several possibilities for the consistent overestimation error: attentional sharing between motor action and monitoring passage of time, or potential anchoring to one second as the test duration was 800 ms. We further distinguish these in Experiment 2.

### Experiment 2

To rule out possible anchoring to one second, we tested two durations, one sub-second (800 ms) and one super-second (1200 ms) in Experiment 2. As we demonstrated that the accuracy feedback was a crucial factor influencing reproduction, we compared the presence and absence of the accuracy feedback as well in Experiment 2. Introducing both the short (800 ms) and long (1200 ms) durations permits us to distinguish two possible accounts: the attentional sharing account and switching accounts. If the general overestimation is a result of attentional switch, we expect comparable overestimation for both short and long durations. In contrast, if overestimation is driven by attentional sharing, we expected overestimation would be larger for the long compared to the short duration. However, according to the scalar property in time perception (Gibbon, 1977; Gibbon & Church, 1990; Ren et al., 2021; Shi et al., 2022), errors measured in ratio should be comparable for the two durations.

### Method

#### Participants and apparatus

Experiment 2 also recruited 20 volunteers to participate (21-33 years old, average of 25.6, 12 females and 8 males). The recruitment and compensation were identical to those in Experiment 1, and the experiment took place in the same cabin and under the same settings as in Experiment 1. The experimental program was also developed using the “Psychopy” package (Peirce et al., 2019) in PyCharm.

#### Stimuli and Procedure

The stimulus in Experiment 2 was the same Gabor patches as in Experiment 1. In addition to the 800ms target duration in Experiment 1, an additional 1200ms target duration was introduced in Experiment 2. These two target durations were tested in two separate sessions to avoid any central tendency effect. During the reproduction process, participants were presented with the same Gabor patch as visual feedback. Unlike Experiment 1, Experiment 2 did not involve any shortened visual feedback. Instead, the output of the action (i.e., visual feedback) was synchronized with the reproduction, meaning it began and ended simultaneously with the reproduction onset and offset. In addition, Experiment 2 compared the presence and absence of the accuracy feedback in the test phase.

Experiment 2 employed the same reproduction task as in Experiment 1, investigating how attention was allocated in reproduction with differentiated durations and the presence of accuracy feedback. Two sessions with different durations (800 and 1200ms) were tested in a succession with a 5 min break in between. The order of the two sessions were counterbalanced among participants. Each session consisted of two phases: an adaptation phase and a test phase. Like in Experiment 1, the adaptation phase included 60 mandatory trials, with an accuracy feedback by the end of each trial. The same criteria as in Experiment 1 were employed to determine the accuracy range and the requirements for passing the adaptation phase. The subsequent test phase consisted of four sub-blocks, each with 25 trials. Regarding the effect of the accuracy feedback, two types of test block were designed: those with accuracy feedback and those without. Specifically, the first and third test blocks supplied accuracy feedback at the end of each trial, while the second and fourth test blocks did not provide any accuracy feedback. This design formed three types of block within each session (800 and 1200 ms): Adaptation/Feedback, Test/Feedback, and Test/No-Feedback blocks.

## Result

Figure 4 shows the mean reproduction durations for two separate durations (800 ms and 1200 ms) in three types of blocks: adaptation, test with accuracy feedback, and test without accuracy feedback. By visual inspection, reproduction was overestimated in the test without accuracy feedback blocks for both durations. The mean reproduced duration (± SE) were 800 (± 11), 806 (± 10), and 910 (± 22) ms for the 800 ms duration, and 1140 (± 10), 1162 (± 14), and 1291 (± 24) ms for the 1200 ms duration, for the adaptation, test with accuracy feedback, and test without accuracy feedback blocks, respectively.

**Figure 4.**
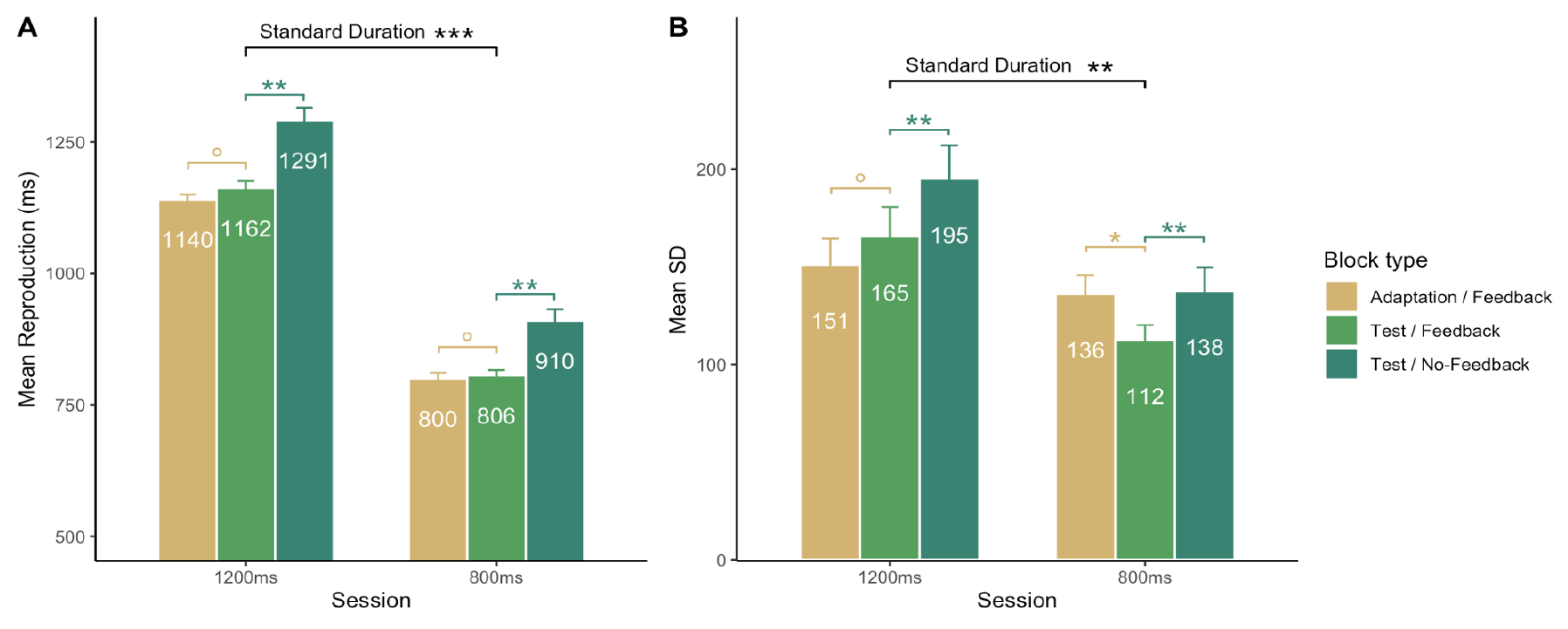
General mean reproduction duration (**A**) and mean standard deviation (SD) (**B**) in Experiment 2, for each session (800 / 1200 ms) and block type (Adaptation/Feedback, Test/Feedback, and Test/No-Feedback). The asterisks mark the significant differences between conditions (p > 0.05: °; p < 0.05: *; p < 0.01:* * ; p < 0.001:* * *).

A 2 (Standard-Duration: 800ms, 1200ms) × 2 (Phase: Adaptation/Feedback, Test/Feedback) ANOVA was conducted comparing the adaptation and test phase with different standard duration. The reproduction difference between 800ms and 1200ms standard duration was significant, (*F*(1, 19) = 1420.06, *p* < .001,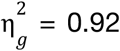), with consistent longer reproduction duration in the 1200ms session compared to in the 800ms session. Neither the main effect of Phase (*F*(1, 19) = 3.24, *p* = .088, 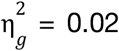) nor the interaction effect between the two main factors was significant (*F*(1, 19) = 0.85, *p* = .368,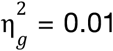), evidencing comparable reproduction in adaptation and test phase when accuracy feedback present in both sessions. A follow-up 2 (Standard-Duration: 800ms, 1200ms) × 2 (Accuracy-Feedback: Test/Feedback, Test/No-Feedback) ANOVA were conducted focusing on the Test phase with accuracy feedback presence and absence. It revealed significant main effect of both Standard-Duration (*F*(1, 19) = 792.83, *p* < .001,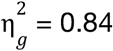) and Accuracy-Feedback (*F*(1, 19) = 48.42, *p* < .001, 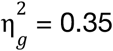), and non-significant interaction effect (*F*(1, 19) = 1.05, *p* = .319, 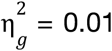). These results are indicative of a general tendency of over-reproduction when the accuracy feedback is absent, independently to the standard duration of reproduction.

The mean standard deviation (SD ± SE) were 136 (± 10), 112 (± 8), and 138 (± 12) ms in the 800 ms session, and 151 (± 13), 165 (± 15), and 195 (± 17) ms in the 1200 ms session (Figure 4), for the Adaptation/Feedback, Test/Feedback, and Test/No-Feedback blocks, respectively. The ANOVA with factor of Standard-Duration and Phase revealed significant main effect for Standard-Duration (*F*(1, 19) = 10.87, *p* = .004, 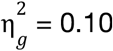) and non-significant main effect of Phase (*F*(1, 19) = 0.28, *p* = .605, 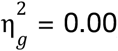). The interaction effect (*F*(1, 19) = 5.26, *p* = .033, 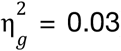) was significant. The interaction was largely due to the relatively large SD in the Adaptation phase of the 800ms session. Further post-hoc comparisons between the Adaptation/Feedback and Test/Feedback blocks revealed a significant difference in the 800ms session (*t*(19) = 2.84, *p* = .011, *BF* = 4.909) but not in the 1200ms session (*t*(19) = -1.00, *p* = .331, *BF* = 0.360). The large SD was mainly induced by the deviation in the early trials in the adaptation phase. The follow-up ANOVA with a specific focus on influence of accuracy feedback in the test phase revealed significant main effect for both Standard-Duration (*F*(1, 19) = 23.38, *p* < .001, 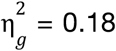) and Accuracy-Feedback (*F*(1, 19) = 13.14, *p* = .002, 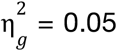). The interaction effect between two main factors (*F*(1, 19) = 0.08, *p* = .783, 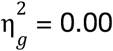) was not significant. It revealed the sensitivity of reproduction performance on the continuous accuracy feedback calibration, suggesting a larger spreading reproduction when the accuracy feedback is absent and higher consistency when present. In summary, the results of Experiment 2 demonstrated that without immediate accuracy feedback calibration, the reproduction was overestimated.

### Constant Overestimation Ratio

The key question to resolve in distinguishing between the two hypothesized attentional mechanisms: attentional sharing and switching -relates to whether overestimation is absolute or ratio-based. As shown in the above results, there was no significant difference between reproduced durations between the adaptation and test with the presence of accuracy feedback. However, a consistent overestimation occurred between the blocks with and without accuracy feedback, for both 800ms and 1200ms sessions. The average overestimation in the 800ms condition was 110 ms, representing 13.8% of the reproduced duration in the feedback blocks. Similarly, in the 1200ms condition, the overestimation was on average 150 ms and accounted for 13.2%. The absolute errors were significantly different between two sessions (*t*(19) = 2.14, *p* = .046, *BF* = 0.691), however, the ratio-based errors were comparable, *t*(19) = 0.28, *p* = .781, *BF* = 0.314.

Recall Experiment 1 had the absolute mean error of 109 ms, we compared the ratio-based errors between two experiments. The average ratio-based errors in the absence of accuracy feedback were 13.7% and 13.5% in Experiments 1 and 2, respectively. A Welch ’s t-test revealed they were comparable, *t*(46) = 0.09, *p* = .929, *BF* = 0.276, suggesting the general overestimation is comparable across different durations and experiments on ratio-based.

## Discussion

In this study, we explored the underlying mechanisms responsible for consistent overestimation in time reproduction, focusing on the influence of both sensory and accuracy feedback during the reproduction process. Experiment 1 involved the shortening of a 150-ms action output (visual feedback), demonstrating that participants can calibrate their reproduction to motorically pressed duration for both shortened and synchronized action outputs if accuracy feedback was provided. When accuracy feedback was absent, however, reproduction progressively drifted back to overestimation in both scenarios. Interestingly, the difference between the shortened and synchronized conditions was consistently maintained across 20 trials in the mini-block. To rule out this drifting was caused by the anchoring effect (where subseconds and supper-seconds tend to anchor to a full second), we compared reproductions for both subsecond (800 ms) and supersecond (1200 ms) durations in Experiment 2. Considering the pivotal role of accuracy feedback in Experiment 1, we also compared reproductions with and without accuracy feedback in Experiment 2. The results indicated a consistent overestimation across both durations, effectively negating the possibility of an anchoring effect. Furthermore, the overestimation, measured in a ratio basis, turned out to be comparable across the different durations and experiments.

In real-world situations, our actions and their effects are typically synchronized. When you press a light switch, the light comes on instantaneously. We only account for a delay when it is noticeable or when we have prior knowledge of it, such as knowing that hot water from a faucet will take a few seconds to flow. This implicit understanding of synchronized actions and effects often shapes how we act. Research has demonstrated that when a delay is subtly introduced between an action and its effect, it can profoundly influence our perception after we have adapted to it (Cai et al., 2012; Cunningham et al., 2001; Ganzenmüller et al., 2012; Shi et al., 2008; Stetson et al., 2006). For instance, when a 100 ms delay was injected between a button press and a subsequent flash, Cai et al. (2012) found that, following adaptation, a synchronized flash with the button press was perceived to have occurred earlier. This means that observers incorporated the 100 ms delay into the action-effect loop and recalibrated their action-effect prior. In Experiment 1 of our study, we also injected a 150 ms onset delay between the initiation of reproduction and its visual feedback. The focus on accuracy feedback was to direct observers ’ attention to their motor timing rather than visual feedback. However, we observed a consistent adaptation effect in the test phase: reproduction during the session with shortened visual feedback was 55 ms shorter compared to the session with synchronized feedback. This reduction in reproduction duration remained stable even without accuracy feedback and during reproduction reverting back the overestimation regime. These findings suggest that a delayed action output can recalibrate the internal prior of perceived duration. The effect was evident and sustained at least through the tested mini-block.

Another intriguing observation from Experiment 1 is the gradual return to an overestimation zone in both sessions. The reversion occurred during the absence of accuracy feedback. This progressively shift was unlikely to occur at the encoding stage, as the visual duration remained unchanged throughout the entire experiment, across two sessions. Rather, the most plausible explanation is that this general overestimation was a trait of the reproduction process itself, and it was constantly recalibrated when the accuracy feedback was provided. The reproduction in the synchronized visual feedback condition stabilized at 927 ms without accuracy feedback, resulting in an over-reproduction of 127 ms when compared to the physical duration of 800 ms. By observing this similar progressively reversion process, we can infer that a comparable overestimation likely remained in the shortened visual feedback condition, even after the prior had been updated during the adaptation to the shortened visual feedback.

So what could be the key factor causing such general overestimation observed during duration reproduction? As we have shown that reproduction with visual feedback cannot be solely attributed to motoric timing. Experiment 1, using shortened visual feedback adaptation, clearly illustrates that both the motor action and its effect (visual feedback) contribute to this general bias. In the introduction, we reviewed two potential explanations for this bias: attentional sharing and attentional switching (Fortin & Rousseau, 1998; Ganzenmüller et al., 2012; Zakay & Block, 1996). The attentional sharing hypothesis (Fortin, 2003; Fortin & Rousseau, 1998; Lejeune, 1998) suggests that attention is divided between action and timing process, resulting in an attentional lapse in monitoring the passage of time. According to the classic pacemaker-switch-accumulator clock model (Gibbon, 1977; Gibbon et al., 1984; Treisman, 1963) and attention gate theory (Zakay, 1989; Zakay & Block, 1996), attentional lapse causes fewer timing ‘ticks ’ reaching the accumulator. The longer the duration is, the more ‘ticks ’ that might be lost. On the other hand, the attentional switch assumes that switching latencies are affected by the task, such as switching from initiating the motor action to beginning to monitor the passage of time. And this switch effect only takes place at the opening and closing of the switch in the pacemaker-switch-accumulator model. Thus, the attentional switch hypothesis predicts that the overestimation should be consistent across different durations, as each duration is only influenced at its onset and the offset. It is worth noting that this interpretation of attentional switching can be seen as a special case of attentional sharing, where impacts occur only at the beginning and end. The interpretation of the ‘ticket ’ loss caused by attentional lapse is closely related to the flickering switch account (Penney et al., 2000). According to this account, observers must maintain constant attention to a stimulus to keep the connection between the pacemaker and the accumulator. Without such attention, the switch may open spontaneously, reducing the number of pulses accumulated (Wearden et al., 2010; see also Wearden, Goodson, et al., 2007). In essence, both the flickering switch and attentional sharing accounts agree that attention is a pivotal factor for the general overestimation in duration reproduction.

In the present study, we observed the increased overestimation as the duration lengthened, which effectively dismisses the pure attentional switch hypothesis. Interestingly, across different sessions in both experiments, we observed that overestimates, calculated on ratio basis, were comparable, at 13.3% and 13.5% in Experiments 1 and 2, in the absence of accuracy feedback. Thus, our data seem to support the attentional sharing hypothesis (Fortin, 2003; Fortin & Rousseau, 1998; Lejeune, 1998) or the flickering switch account (Penney et al., 2000). However, we cannot entirely rule out that the possibility of the pacemaker speed of the temporal accumulation might slow down (Wearden et al., 2010; Wearden, Goodson, et al., 2007). This slowing down of pacemaker speed would mimic the prediction of the attentional sharing account (Fortin, 2003; Fortin & Rousseau, 1998; Lejeune, 1998) -a general overestimation is proportionate to the test duration. The present study, unfortunately, is unable to fully differentiate these two alternative explanations. Future research using extremely short or long durations (Wearden, Norton, et al., 2007) could further help untangle these two accounts.

However, insights might be drawn from a recent study (Ren et al., 2021). Ren and colleagues (2021) carried out a duration reproduction study across an extensive range, from 300 ms to 16 seconds. They observed consistent overestimation when durations were tested in blocks (separated into subsecond, seconds, and super-seconds categories to reduce the central tendency effect), except for the extreme long duration 16 seconds, where an underestimate was detected. Their findings implies that the general overestimation bias does not uniformly apply across all durations but is mainly confined to the sub-second and second ranges. For extremely long durations, the general bias appears to shift towards an underestimation. Such underestimation might partially be ascribed to the central tendency effect. As our day-to-day activities requiring critical actions usually fall within the subsecond and second ranges, this creates an implicit prior for action-related events in these time frames. When tasked with reproducing an extraordinarily long duration, this implicit prior likely influences the reproduction outcome. However, attentional lapse could also contribute to this underestimation. In the case of prolonged durations (such as 16 seconds), both attentional lapse and memory decay might affect the encoding and reproduction phases, and with significant impact of the former, resulting in an underestimation. Such underestimation phenomenon is challenging to explain solely with the slow-down pacemaker speed account, as it would predict the opposite outcome -overestimation.

In conclusion, here we observed a consistent overestimation in the reproduction of durations around a second, in the absence of accuracy feedback. The overestimation was approximately 13.5% on ratio basis across different durations and sessions, which was not influenced by shortened visual feedback. We propose that this consistent overestimation is a result of attentional sharing between action and monitoring of the passage of time during the reproduction process. Further research is needed to further disentangle roles of the pacemaker speed and attention in temporal reproduction.

